# SPEAR: a proteomics approach for simultaneous protein expression and redox analysis

**DOI:** 10.1101/2021.06.17.448798

**Authors:** Shani Doron, Nardy Lampl, Alon Savidor, Corine Katina, Alexandra Gabashvili, Yishai Levin, Shilo Rosenwasser

## Abstract

Oxidation and reduction of protein cysteinyl thiols serve as molecular switches, which is considered the most central mechanism for redox regulation of biological processes, altering protein structure, biochemical activity, subcellular localization, and binding affinity. Redox proteomics allows for the global identification of redox-modified cysteine (Cys) sites and quantification of their oxidation/reduction responses, serving as a hypothesis-generating platform to stimulate redox biology mechanistic research. Here, we developed Simultaneous Protein Expression and Redox (SPEAR) analysis, a new redox-proteomics approach based on differential labeling of oxidized and reduced cysteines with light and heavy isotopic forms of commercially available isotopically-labeled N-ethylmaleimide (NEM). The presented method does not require enrichment for labeled peptides, thus enabling simultaneous quantification of Cys oxidation state and protein abundance. Using SPEAR, we were able to quantify the in-vivo oxidation state of thousands of cysteines across the *Arabidopsis* proteome under steady-state and oxidative stress conditions. Functional assignment of the identified redox-sensitive proteins demonstrated the widespread effect of oxidative conditions on various cellular functions and highlighted the enrichment of chloroplast-targeted proteins. SPEAR provides a simple, straightforward, and cost-effective means of studying redox proteome dynamics. The presented data provide a global quantitative view of cysteine oxidation of well-known redox-regulated active sites and many novel redox-sensitive sites whose role in plant acclimation to stress conditions remains to be further explored.

## Introduction

As sessile organisms that grow under dynamic environmental conditions, plants must constantly sense, respond, and adapt to fluctuations in their environment. An increase in reactive oxygen species (ROS) production, emanating from oxygen-based metabolic pathways, such as photosynthesis, photorespiration and oxidative phosphorylation, has been shown to play a signaling role in plant adaption to stress conditions (Foyer and Noctor, 2005; Gadjev et al., 2006; Halliwell, 2006; Suzuki et al., 2012). Oxidative stress is a unique physiological state characterized by modulation of gene expression patterns (Gadjev et al., 2006; Rosenwasser et al., 2013; Willems et al., 2016) and metabolic activities involving ROS as a secondary messenger within complex signaling networks (Laloi et al., 2004; Halliwell, 2006). Specifically, hydrogen peroxide (H_2_O_2_), one of the most studied ROS, acts as an essential messenger molecule mediating diverse cellular processes. H_2_O_2_ or redox changes can be sensed by monitoring the reversible oxidation of cysteine (Cys) thiol (−SH) groups to sulfenic acid (−SOH) (Winterbourn and Hampton, 2008; Petrov and Van Breusegem, 2012). The highly unstable sulfenic acid can further be oxidized by ROS, forming sulfinic acid (−SO_2_H) and sulfonic acid (−SO_3_H), or react with other cysteine residues, forming disulfide bonds (−SSR) (Yang et al., 2016). Oxidation of these reactive cysteines acts as a molecular switch, which is considered the most central mechanism for redox regulation of biological processes, altering protein structure, biochemical activity, subcellular localization and binding affinity (Waszczak et al., 2015). For instance, thiol-based signal transduction, which was first described as a regulator of chloroplast stroma activities during light-driven electron flux (Buchanan and Balmer, 2005), was found to be particularly important in photosynthetic organisms that have to rapidly respond to fluctuating light intensities and multiple environmental cues (Dietz and Pfannschmidt, 2011; Scheibe and Dietz, 2012). Thus, identifying redox-modified proteins and quantifying cysteine reactivity is critical in understanding how plants perceive and respond to transducing redox signals in changing environmental conditions.

Recent advancements in mass spectrometry (MS) technology have led to the development of various proteomics techniques to detect reversibly modified cysteines, which have enabled system-level identification of redox-sensitive proteins. Thioredoxins (Trxs) and glutaredoxins (Grxs)-regulated proteins in plants were thoroughly characterized by applying trapping chromatography using single-cysteine Trx or Grx mutant which forms a stable mixed disulfide bond with target proteins (Balmer et al., 2004; Rouhier et al., 2005; Yoshida et al., 2013; Buchanan, 2016). Such methods allowed the identification of numerous redox-regulated proteins and demonstrated the prevalence of redox regulation across plant metabolic pathways and molecular processes. However, a more in-depth understanding of redox dynamics requires *in-vivo* quantification of the redox-modified site under diverse physiological conditions, which is not enabled by monothiol mutant-based trapping approaches.

Several gel-free, site-specific, and quantitative redox approaches have been developed to identify redox-sensitive cysteine sites (Lennicke et al., 2016; Yu et al., 2020). For example, the OxiTRAQ uses the biotin-switch method to differentially label thiol groups, which can then be identified and quantified using the iTRAQ method, as applied to suspended *Arabidopsis* cells treated with H_2_O_2_ (Liu et al., 2014). Relative quantitation of reversibly oxidized cysteine-containing peptides using the Cys tandem mass tag (cysTMT) was demonstrated in tomato plants in response to *Pseudomonas syringae* (Balmant et al., 2015). When irreversible labeling of Cys using iodoTMT reagents is combined with iTRAQ tags, both changes in protein levels can be quantified and redox-modified Cys in proteins identified (Parker et al., 2015). This hybrid approach, named iodoTMTRAQ, was recently employed to analyze changes in protein redox modifications in suspended *Arabidopsis thaliana* cells in response to bicarbonate (Yin et al., 2017). As most redox proteomics approaches use reducing agents such as DTT or TCEP to indiscriminately reduce all modified cysteines, the exact chemical Cys modification cannot be distinguished. Thus, identifying a precise modification requires either the use of a specific reducing reagent or direct thiol labeling of a specific Cys modification. For example, using a set of S-sulfenylation-specific probes enabled highly comprehensive characterization of S-sulfenylated cysteines in *Arabidopsis* cells (Waszczak et al., 2014; Akter et al., 2015; De Smet et al., 2019; Huang et al., 2019). Most quantitative redox proteomics analyses detect either the reduced or the oxidized state of a given thiol, presenting fold-change values between control and treatment by comparing the incorporation of an alkylating probe in different samples (Mock and Dietz, 2016). In contrast, redox proteomics methods that are based on differential labeling of reduced and oxidized thiols with isotopically light and heavy forms of thiol-alkylating agents, such as the OxiCAT or iodoacetyl tandem mass tag (iodoTMT), enable precise measurement of the percent of cysteine oxidation (Leichert et al., 2008; Brandes et al., 2011; Nietzel et al., 2020). The later value is of high importance to identify Cys that their redox modulation has significant physiological effects. Recently, OxiCAT provided a comprehensive view on the diatom *Phaeodactylum tricornutum* redoxome and insights into its evolution (Rosenwasser et al., 2014; Woehle et al., 2017). Notably, as OxiCAT and iodoTMT requires the enrichment of cysteine-labeled peptides, data regarding protein expression is lost during sample processing and cannot be analyzed simultaneously with cysteine redox changes. Also, the intensive manual work involved in peptide purification by affinity chromatography and the cost of chemicals required for Cys trapping limits the number of samples that can be analyzed in a single experiment.

Two recently developed redox proteomics approaches allow determining the percent of oxidation of identified Cys-containing peptides and relative quantification of protein abundance in the same experiments (Anjo et al., 2019; Xiao et al., 2020). Deep coverage and quantification of the tissue-specific mouse Cys proteome were achieved using IAM-based cysteine-reactive phosphate tags coupled with TMT isobaric reagents (Xiao et al., 2020). Using commonly available alkylating reagents, oxSWATH combines label-free protein expression profiling with differential alkylation to quantify relative redox changes on a global scale (Anjo et al., 2019). These protocols are based on splitting the original sample into two or three identical sub-samples that undergo complementary differential alkylation labeling procedures and have not been adopted to plant tissue.

Here, we developed a new redox-proteomics approach that is based on differential labeling of oxidized and reduced cysteines with commercially available light (d0) and heavy (d5) isotopic forms of the common thiol-alkylating reagent N-ethylmaleimide (NEM). By mixing samples in which reduced cysteines were labeled with d0-NEM or d5-NEM, peptide fractionation using high pH reversed-phase chromatography, online nanoflow liquid chromatography (nanoAcquity), high mass accuracy, tandem mass spectrometry (Q-Exactive HF), and a next-generation search engine, we were able to quantify the *in-vivo* oxidation state of thousands of cysteines across the *Arabidopsis* proteome under steady-state and oxidative stress conditions. The presented data provide a global quantitative view of cysteine oxidation in photosensitizing plant leaves.

## Results and Discussion

### Intracellular response to hydrogen peroxide

We aimed to detect and quantify reactive thiols on a system level, under steady-state conditions and following exposure to physiologically relevant hydrogen peroxide (H_2_O_2_) concentrations. To examine the intracellular response to H_2_O_2_, we measured the oxidation state of chloroplast 2-Cys peroxiredoxin (2-Cys-Prx), as previously demonstrated for isolated chloroplasts (Muthuramalingam et al., 2013). 2-Cys-Prx can serve as a sensor for intracellular oxidation state, reflected in its oligomeric form. The oxidation state of 2-Cys-Prx was determined by measuring the relative amounts of the dimeric (oxidative) versus monomeric (reduced or hyperoxidized) forms on non-reducing SDS-PAGE using a 2-Cys-Prx-specific antibody. As shown in Fig. 1a&b, a major increase in the relative abundance of the 2-Cys-Prx oxidized states (out of total 2-Cys-Prx) from 56% to 88% and 80% was achieved following the application of 10mM and 50mM H_2_O_2_, respectively. The observed monomeric form may also represent the hyperoxidized 2-Cys-Prx, the higher oxidation state of 2-Cys-Prx catalytic site sulfonic (−SO_3_^−^) derivative. A high level of hyperoxidized 2-Cys-Prx may be an indication of deleterious H_2_O_2_ concentrations and activities aimed at protecting cells against oxidative damage by maintaining the reduced state of thioredoxins and increasing signaling via 2-Cys-Prx. (Veal et al., 2018). Thus, an accumulation of the hyperoxidized 2-Cys-Prx was measured using an antibody raised specifically against this irreversible (−SO_3_^−^) hyperoxidized state (Fig. 1a low panel &b). Only a minor and moderate increase in hyperoxidation 2-Cys-Prx was observed following treatment with 10mM and 50mMH_2_O_2_, respectively, while a significant increase was detected following treatment with 100mM H_2_O_2_.

**Figure 1.**
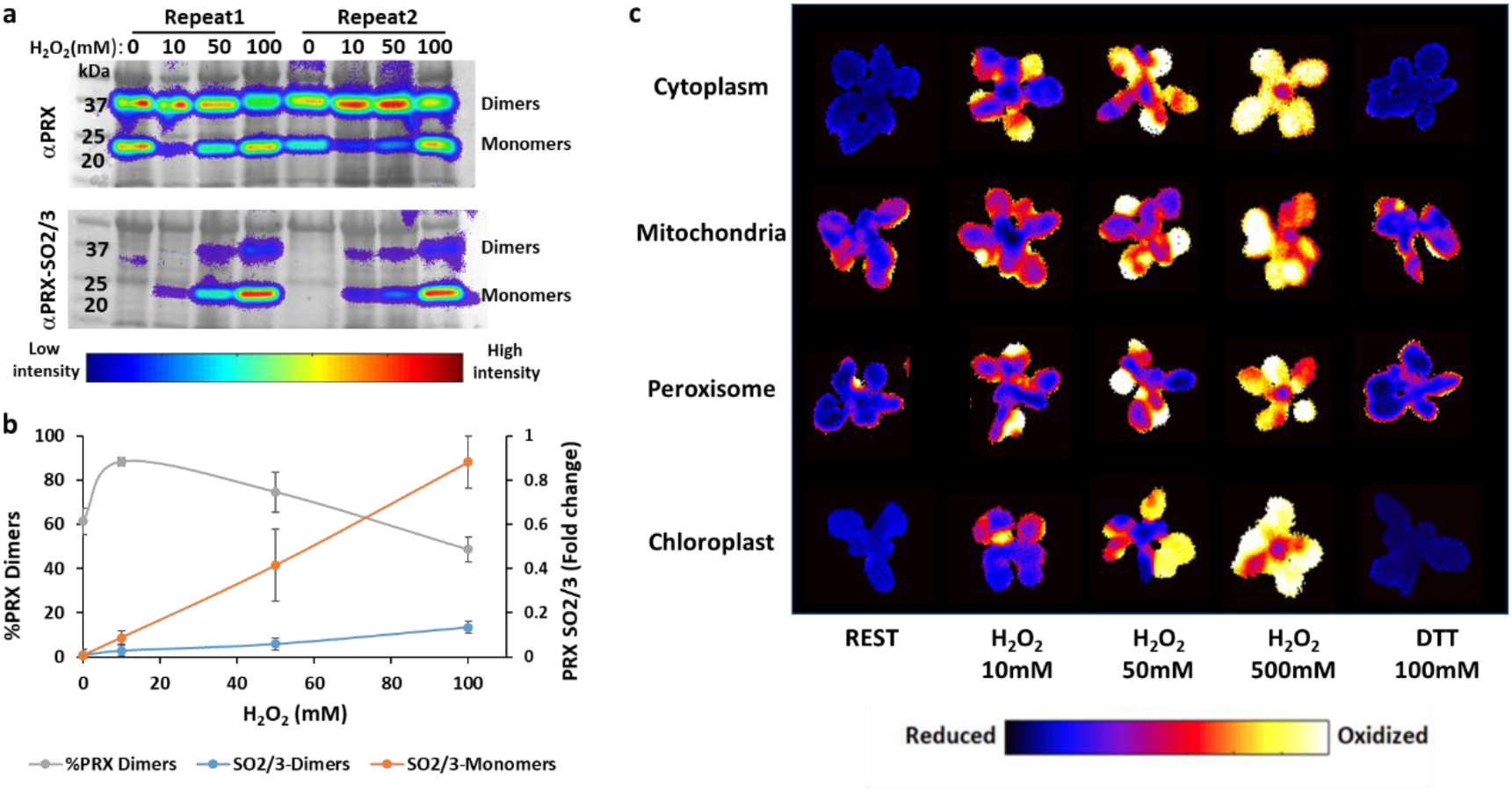
Dose-dependent response of chloroplastic 2-Cys peroxiredoxin and organelle-targeted roGFP2 to increasing concentrations of H_2_O_2_. **(a)** Immunoblot showing the oxidized state of 2-Cys peroxiredoxin (upper panel) and the hyperoxidized forms of peroxiredoxin −SO_3_ (lower panel) in *Arabidopsis* plants after exposure to increasing concentrations of H_2_O_2_ (10-100 mM) for 20min. Total protein was extracted using TCA and free thiols were blocked with N-ethylmaleimide (NEM). **(b)** Quantification of the relative abundance of oxidized 2-Cys peroxiredoxin (dimer) and hyperoxidized peroxiredoxin forms presented in **a**. The relative abundance of the hyperoxidized form was quantified as a percentage of the intensity measured following treatment with 0-100mM H_2_O_2_. Values represent the mean ±SD of two biological replicates. **(c)** Whole-plant ratiometric imaging of *Arabidopsis* plants expressing roGFP2 in the cytoplasm, mitochondria, peroxisome, or chloroplasts, and subjected to H_2_O_2_ and DTT treatments.

To further check whether the exogenously applied peroxide significantly affects the reactive thiols in other subcellular localizations, we examined the oxidation state of the redox-specific *in-vivo* sensor, roGFP2, targeted to specific subcellular compartments in response to exogenous application of H_2_O_2_ (Fig. 1c). Whole-plant fluorescence imaging analysis showed partial oxidation of the probe in the chloroplast, mitochondria, peroxisome and cytoplasm following treatment with 10mM and 50mM H_2_O_2_, while full oxidation was only achieved following treatment with 500mM H_2_O_2_ (Fig. 1c). Thus, concentrations of 10mM and 50mM H_2_O_2_ were further used to represent physiological H_2_O_2_ concentrations that affect proteins in diverse subcellular localizations.

### Global quantification of Cys oxidation

To comprehensively characterize the cellular responses of *Arabidopsis* leaves to the early phase of oxidative stress, we aimed to detect and quantify reactive thiols on a system level, under steady-state conditions and following exposure to physiological hydrogen peroxide (H_2_O_2_) concentrations. Our approach for global quantification of the Cys redox state, termed SPEAR (simultaneous protein expression and redox analysis), is based on rapid acidification of plant extracts with trichloroacetic acid (TCA) to preserve redox status of thiol groups, followed by differential labeling with isotopically light (d0) (reduced cysteines), and heavy (d5) (oxidized cysteines) forms of NEM, as demonstrated by (McDonagh et al., 2014, Fig. 2a). The identity and abundance of the reduced and the oxidized forms of specific peptides can be determined based on 5 Da mass difference between the light and heavy peptides. Trypsin-digested peptides were directly analyzed by LC-MS/MS proteomics without enrichment of cysteine-containing peptides, thus also enabling protein level quantification, since the data includes non-modified peptides of these proteins.

**Figure 2:**
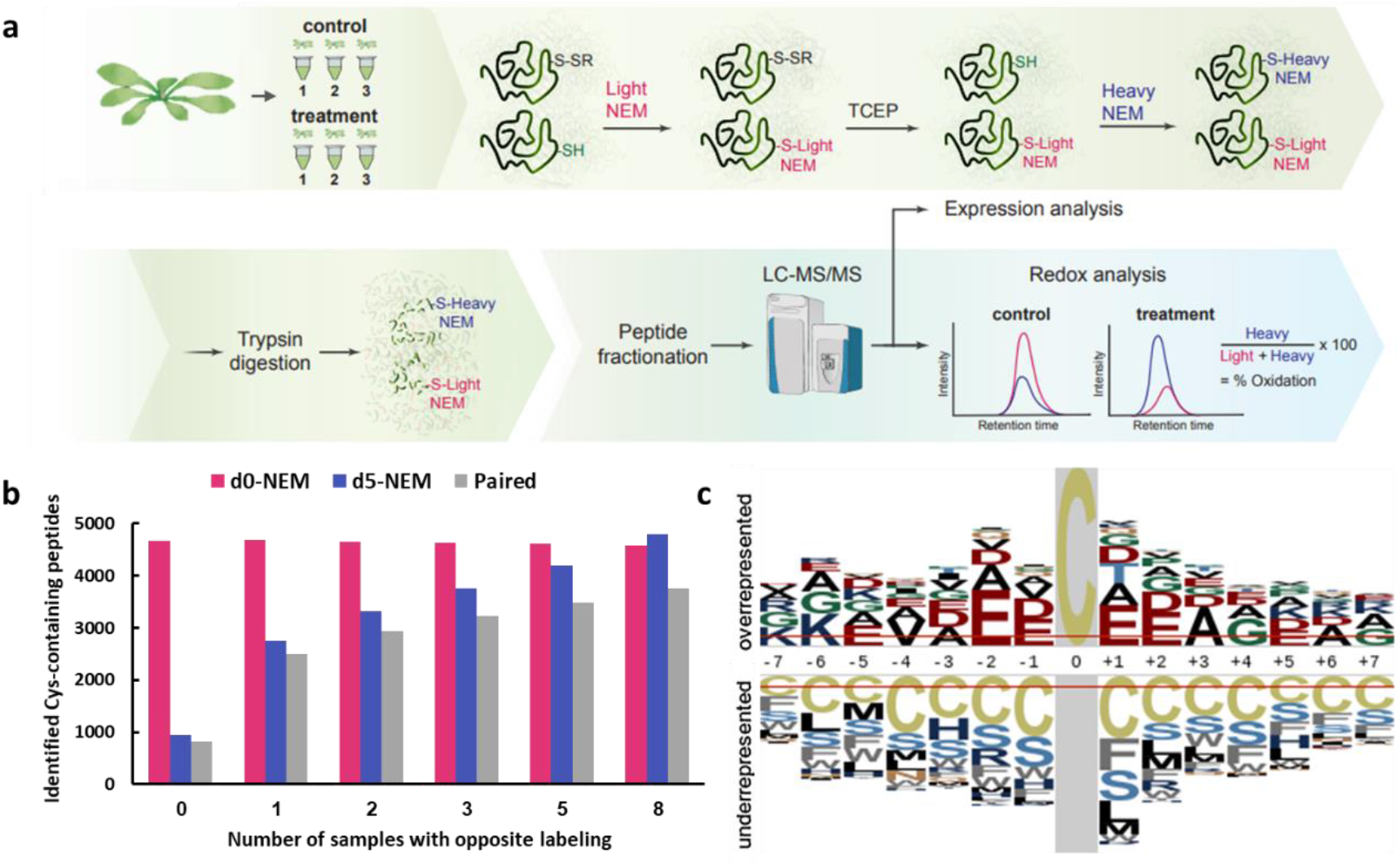
Simultaneous quantification of Cys oxidation state and protein expression levels using the SPEAR approach. **(a)** Schematic representation of the NEM-based differential alkylation method used to evaluate Cys degree of oxidation and protein abundance. S-SR and SH signify an oxidized and a reduced Cys of a represented redox sensitive protein, respectively. The abundance of the reduced and the oxidized forms of a specific peptide can be distinguished based on the 5 Da mass difference between the light- and heavy-labeled peptides. As no enrichment and purification steps were included, data regarding protein expression was maintained and extracted using common label-free protein quantification approaches. **(b)** The number of identified peptide pairs (labeled with d0 and d5 NEM) was greatly enhanced by the co-analysis of peptides labeled as described in A and peptides in which reduced Cys were labeled with d5 NEM (opposite labeling). The increase in the number of identified d0-labeled, d5 labeled peptides and pairs due to the addition of samples with opposite labeling using only the G-PTM-D approach is presented. **(c)** The consensus motif of NEM-labeled peptides demonstrates significant enrichment of aspartic and glutamic acid near the modified Cys. The *Arabidopsis thaliana* proteome was used as the proteomic background population for the calculation of the probabilities. Similar analysis done for non-labeled Cys containing peptides is presented in Supp. Fig. 2.

To expand the total number of identified peptides, sample fractionation using high pH reversed-phase chromatography was carried out, followed by online nanoflow liquid chromatography (nanoAcquity) and high-resolution high mass accuracy, tandem mass spectrometry (Q-Exactive HF). The recently developed MetaMorpheus (MM), a next-generation search engine (Solntsev et al., 2018), was employed to identify and quantify light and heavy NEM-labeled peptides. Quantification was achieved by comparing the area under the curve of the light and heavy-labeled peptides at the intact peptide level (MS1). Importantly, unlike differential labeling with carbon isotopes (as used in OxiCAT), in which the light and heavy forms of the same peptide elute at the same time, the difference in hydrophobicity between hydrogen and deuterium of the d0-NEM and d5-NEM peptides results in unequal retention times (RT) (Fig. S1). For this reason, the RT data cannot be used in matching light and heavy peptides, and identification by MS/MS of each form is required.

To test the capabilities of this relatively simple and straightforward workflow, plants were treated with 10mM or 50mM H_2_O_2_ for 20 min, as was done in the aforementioned experiments to establish the physiological conditions, and subjected to d0-NEM and d5-NEM labeling of reduced and oxidized Cys thiols, respectively (Fig. 2a). Two biological replicates were performed f each treatment, resulting in six sample and a total of 36 LC-MS/MS runs. Using the G-PTM-D algorithm of MM, 3689 d0-NEM and 518 d5-NEM labeled unique peptides were identified, enabling the quantification of the degree of oxidation of 412 cysteines. In this dataset, 17 cysteines exhibited a significantly increased degree of oxidation under oxidative stress conditions and were identified as redox-sensitive (Table S1).

We recognized that the relatively small number of identified d5-NEM-labeled peptides compared with d0-NEM labeled peptides resulted from the highly reduced state of most thiol groups, hindering the ability to accurately identify, by MS/MS, the fraction of oxidized peptides due to their low abundance. To overcome this limitation, we expanded our approach by analyzing an additional set of samples in which the reduced Cys groups were labeled with d5-NEM (we referred to this as “opposite labeling”), allowing us to precisely extract their m/z values and their specific RT. Then, by applying the MM ‘match between runs’ algorithm, the MS profiles derived from the opposite labeling samples served as markers to locate the corresponding peptides in the core samples labeled with d0-NEM and d5-NEM for the reduced and oxidized Cys states, respectively.

To examine the applicability of this approach, proteins extracted from control, 10mM and 50mM H_2_O_2_-treated plants were labeled with d0-NEM and d5-NEM for the reduced and oxidized Cys states, respectively. An additional set of samples was oppositely labeled, as described above. As shown in Fig. 2b, the number of d5-NEM-labeled peptides in the eight-core samples increased significantly due to their co-analysis with those having reduced Cys labeled with d5-NEM, resulting in the higher number of identified d5/d0 pairs. These results demonstrated that by adding oppositely labeled “marker” samples, the number of identified Cys could significantly increase without purifying and enriching labeled peptides. In total, double labeling with both the heavy and the light NEM reagents was recorded for 5523 peptides, representing 2967 distinct proteins in the Arabidopsis proteome. This was achieved by combining protein search data of G-PTM-D and variable modification approaches provided in the MM platform, demonstrating in-depth analysis of the redox proteome without bias toward only highly abundant proteins. Most (~96%) of the identified Cys-containing peptides were found to be NEM-labeled, pointing to the high efficiency of the labeling procedure. Interestingly, we found that NEM-labeled cysteine residues were in close proximity to a consensus motif significantly enriched with glutamic acid and aspartic acid (Fig. 2c). This motif was not detected in identified Cys-containing peptides that were not labeled with NEM (Fig. S2), suggesting that these acidic amino acids enhance cysteine reactivity to NEM.

While commonly used redox proteomics approaches are semi-quantitative and report the fold-change between specific oxidation states (reduced or oxidized) in control versus treated samples, the ability of SPEAR to measure both the exact percent of modified cysteines (i.e., site occupancy) and protein levels enables to determine whether the oxidation of specific cysteines alone (i.e. without additional transcriptional/translational regulation) is sufficient to elicit a significant physiological consequence. This valuable information regarding redox regulatory nodes is critical when redox networks are assembled for computational simulation to describe basic parameters of the thiol regulatory networks (Gerken et al., 2020). To conclude, we propose combining peptide fractionation and parallel running of samples with regular and reverse labeling to identify high numbers of labeled pairs without necessitating enrichment for labeled peptides, enabling retention of overall protein abundance data.

### The *Arabidopsis* redox proteome

The degree of oxidation was calculated for each cysteine based on the intensity of the light and heavy peptides. The median degree of oxidation of cysteines under steady-state conditions, following low (10mM) and high (50mM) H_2_O_2_ treatments, was 19.85%, 27.45%, and 28.93%, respectively, demonstrating that the majority of the Cys remained highly reduced even under oxidative stress conditions. In total, we found 577 cysteine-containing peptides, representing 525 proteins, that exhibited a significant change in their degree of oxidation (|ΔOX|>10% and *P*-value < 0.05) under oxidative stress conditions (Fig. 3a&b and Table S2). These results, which indicate the specificity of redox signaling, are consistent with earlier redox proteomics surveys, which showed that only a distinct subset of thiols undergoes stable, yet reversible, thiol modification in response to H_2_O_2_ (Brandes et al., 2011; Liu et al., 2014; Rosenwasser et al., 2014; Fu et al., 2017).

**Figure 3:**
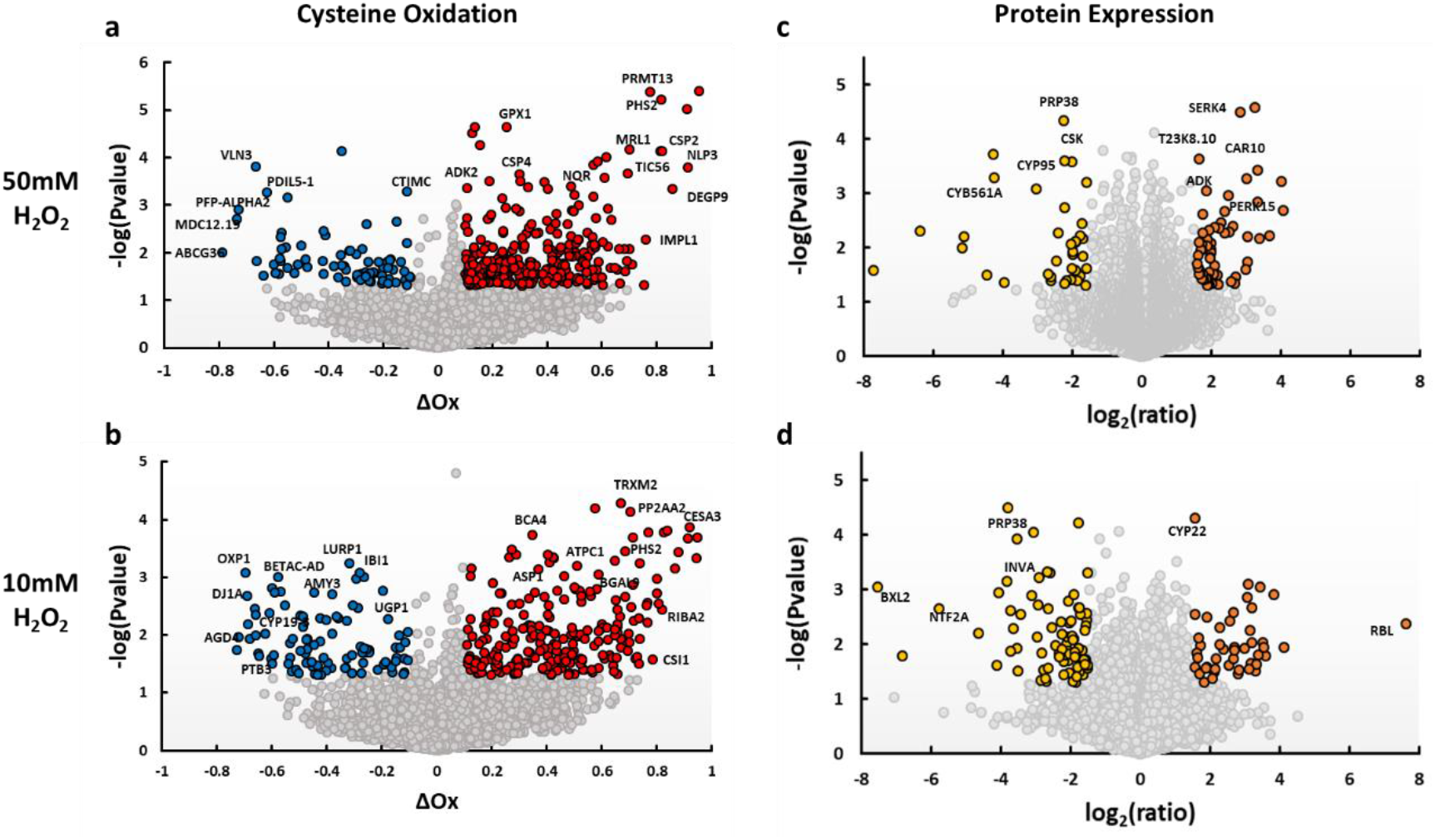
Simultaneous quantification of Cys oxidation state and protein expression levels under oxidative stress conditions. Whole rosettes of 3-week-old *Arabidopsis* plants were soaked in 10mM or 50Mm H_2_0_2_ and subjected to SPEAR analysis. **(a, b)** Volcano plot visualization of the response of 5523 unique cysteine-containing peptides to 10mM **(b)** or 50mM **(a)** H_2_0_2_. Colored dots highlight Cys with a ≥ 10% change in oxidation (ΔOx) and a p-value < 0.05 in H_2_O_2_-treated plants compared to control. **(c, d)** Volcano plot visualization of the expression profile of 7493 proteins following treatment with 10mM **(d)** or 50mM **(c)** H_2_0_2_. Colored dots highlight proteins with an average ≥ 1.5 log2-fold change in H_2_O_2_-treated plants compared to control and a p-value < 0.05.

Examination of protein abundance showed that the level of 127 and 96 proteins was altered in response to a 20-min exposure to 10mM or 50mM H_2_O_2_, respectively (|Log_2_ FC|>1.5, *P*-value < 0.05, Table S3), including upregulation of several protein kinases, such as PROLINE-RICH EXTENSIN-LIKE RECEPTOR KINASE 15 (PERK15). Interestingly, we did not find any overlap between proteins that underwent oxidization and proteins whose expression level changed under oxidative stress, demonstrating the independence of these two regulatory modes. The relatively minor alteration in protein abundance compared with the major cysteine modifications, emphasizes the role of redox regulation in the rapid modulation of protein activity.

SUBA, a tool to investigate proteins subcellular localization in *Arabidopsis* was then used to predict the location of the identified proteins (Hooper et al., 2017). The analysis revealed that redox-sensitive proteins were found in all cell compartments, in agreement with observed roGFP oxidation in all examined subcellular localization (Fig. 1c). It also revealed an over-representation of plastid-targeted proteins (22% of identified proteins and 32% of redox-sensitive proteins, hypergeometric test p = 1.264E^−08^, Fig. 4a&b). Thus, despite the high antioxidant buffer capacity of the chloroplast, its proteome is enriched in redox-sensitive proteins, highlighting the widespread role of redox regulation in chloroplast reactions. Cytosolic proteins were also overrepresented in the redox-sensitive proteins (21% of identified proteins and 28% of redox-sensitive proteins, hypergeometric test p = 4.394E−^05^). Among them were 16 translation-related proteins, in agreement with the ROS sensitivity of the translation machinery recently observed in yeast cells (Topf et al., 2018). GO (Gene Ontology) and KEGG (Kyoto Encyclopedia of Genes and Genomes) assignment of redox-sensitive proteins included the enrichment of redox-related terms, such as peroxiredoxin activity (GO:0051920) and iron-sulfur cluster binding (GO:0051539), as well as the overrepresentation of various metabolic pathways, such as glycolysis (ath00010) and carbon fixation (ath00710) (Fig. 4c).

**Figure 4:**
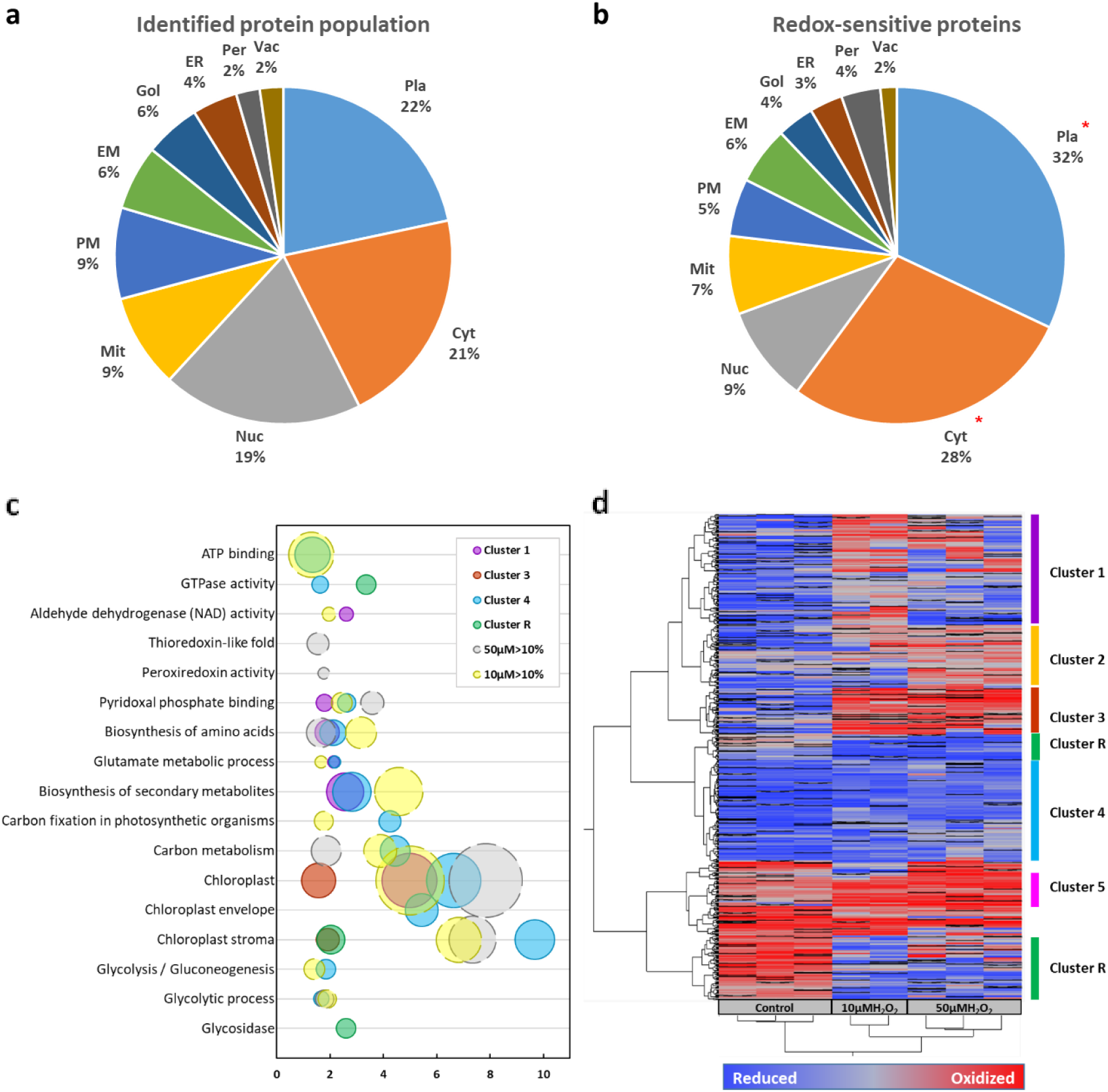
Distribution of redox-sensitive proteins among subcellular compartments and global Cys oxidation profiles of the *Arabidopsis* plant under oxidative stress conditions. **(a,b)** Subcellular distribution of redox-sensitive proteins were assigned using the SUBA tool (Hooper et al., 2017). The percentage of proteins in each compartment from the **(a)** entire identified protein population and **(b)** redox-sensitive proteins is depicted. Distribution of the thiol-oxidized state between steady state and exposure to 10 and 50mM H_2_O_2_ treatment in different subcellular compartments is presented in **Fig. S5**. **(c)** Significantly enriched GO terms and KEGG pathways (hypergeometric test, P < 0.05) in redox-sensitive proteins (change in the oxidation of ≥ 10% and a P-value < 0.05) and clusters as displayed in **(d)**. For a full list of enriched biological pathways in each cluster, see Supplemental Data Set 2. **(d)** Hierarchical clustering of 577 Cys with altered oxidation state in response to 10mM or 50mM H_2_O_2_. Red, high oxidation state; blue, low oxidation state. *hypergeomtric test <0.01. Pla, plastid; Cyt, cytosol; Nuc, nucleus; Mit, mitochondrion; PM, plasma membrane; EM, extracellular matrix; Gol, golgi ; ER, endoplasmic reticulum ; Per, peroxisome ; Vac, vacuole.

Hierarchical clustering of redox-sensitive proteins, based on the degree of oxidation, showed enrichment of several clusters with differential redox sensitivity in specific biological pathways (Fig. 4d). Of note was a cluster of proteins in which their reactive Cys underwent reduction in response to H_2_O_2_ treatment. These counterintuitive results, that were also reported in other redox proteomics analyses (Wang et al., 2012; Liu et al., 2014), demonstrate the transmission of reductive signals under oxidative stress conditions. For example, we detected a reduction of Cys^218^ of the mitochondrial NifU-like protein (ΔOX_L_=−40%, ΔOXH=−57%) and Cys^209^ of chloroplastic phosphoglucomutase (ΔOX_L_=−13%, ΔOX_H_=−11%). In yeast, it has been suggested that hyperoxidation of 2-Cys peroxiredoxin results in preservation of reduced thioredoxin, which then reduces its targets and repairs oxidative damage, providing a rational mechanism for H_2_O_2_ induced reduction of thiols under oxidative stress conditions (Day et al., 2012). A reduction of H_2_O_2_-target proteins in response to peroxiredoxin inactivation is also suggested by the peroxiredoxin redox relay model (Calabrese et al., 2019). However, as shown in Fig. 1a&b, our data indicated only minor hyperoxidation of chloroplastic 2-Cys peroxiredoxin. Thus, the observed reduction of specific protein groups may result from the hyperoxidation of other peroxiredoxins or that even minor levels of 2-Cys peroxiredoxin hyperoxided state are sufficient to induce Cys reduction. Alternatively, besides hyperoxidation, inhibition of metabolic activity that consumes reducing power, such as carbon fixation, may result in a diversion of reducing power to protein targets.

### The functionality of redox-sensitive cysteines

Functional annotation retrieved for all the quantified cysteines from the UniProt knowledgebase (https://www.uniprot.org/, Consortium, 2019) indicated an overrepresentation of active-site cysteines (hypergeometric test, p-value=0.00015, Supp. Table 2) and of redox-sensitive Cys residues involved in disulfide bridges (hypergeometric test, p-value=0.0268, Supp. Table 2) in the redox-sensitive cysteines compared to the all quantified cysteines. Among the observed active-site cysteines were known redox-regulated sites such as glutathione-dependent dehydroascorbate reductase (DHAR) and glutathione peroxidase 1 (GPX1). Other oxidized active site Cys residues included Cys^139^ of ATP synthase gamma chain 1 (ATPC1), Cys^106^ of ferredoxin-dependent glutamate synthase 1 (GLU1), Cys^156^ of Glyceraldehyde-3-phosphate dehydrogenase C subunit (GAPC) and Cys^184^ of plastid rhodanese-like protein 11 (AtStr11). Among the redox-sensitive cysteines involved in disulfide formation were Cys^187^ in magnesium-chelatase subunit ChlI-2 (CHLI2), which has been shown to form a disulfide with Cys^96^ under oxidative conditions (Ikegami et al., 2007), Cys^217^ in NADPH-dependent thioredoxin reductase 3 (NTRC), which forms a disulfide with Cys^220^ and Cys^157^ of glucose-6-phosphate 1-dehydrogenase 1 (G6PDH1), which is involved in redox modulation of the oxidative pentose-phosphate pathway (Wenderoth et al., 1997).

Moreover, twelve reactive sites were found to be involved in zinc (9 sites) and iron-sulfur (3 sites) cluster coordination. Oxidation of these cysteines can lead to metal release, resulting in altered protein conformation and function (D’Autréaux and Toledano, 2007). We also identified and quantified the redox sensitivity of Cys^139^ (ΔOX_H_=38%), which is involved in the coordination of zinc in PSA2 – a thylakoid member of the DnaJ-like zinc finger domain protein family that affects light acclimation and chloroplast development by regulating the biogenesis of photosystem I (PSI) (Fristedt et al., 2014). Oxidation of these Cys residues under oxidative stress conditions can serve as a mechanism that slows down photosynthesis under stress conditions. Cys^644^ (ΔOX_L_=66%) involved in the coordination of iron-sulfur cluster was found in an oxidized state in 4-hydroxy-3-methylbut-2-en-1-yl diphosphate synthase (ISPG), which is involved in the plastid non-mevalonate pathway for isoprenoid biosynthesis. As the electrons required for ISPG activity are directly provided by the electron flow from photosynthesis via ferredoxin, oxidation of these Cys residues may regulate the electron flux through this pathway.

### Quantification of redox reactivity in redox sensors and regulators

An increase in the abundance of the oxidative state of many known redox sensors and regulators was detected following exposure to hydrogen peroxide, reinforcing the validity of the redox proteomics approach and its high sensitivity in capturing *in vivo* states of thiol proteins (Fig. 5a). For example, the extracted ion chromatograms illustrated the redox sensitivity of the Cys^111^ (ΔOX_H_ =16%) of the catalytic site of peroxiredoxin Q (PRXQ) (Fig. 5b). This protein was shown to attach to the thylakoid membrane and was suggested to play a role in protecting photosystem II against hydrogen peroxide (Lamkemeyer et al., 2006). Major oxidation was also measured in the catalytic site of peroxiredoxin-2E (PRXIIE, Cys^121^, ΔOX_H_ =59%), which has been implicated in light acclimation (Rouhier and Jacquot, 2005) and subjected to S-Nitrosylation (Romero-Puertas et al., 2007). Oxidation of the resolving thiol, Cys^241^, of 2-Cys-Prx (BAS1, ΔOX_L_ =23%, ΔOX_H_ =24%, Fig. S3 and Supp. Table 2), was also detected, confirming the shift toward its oxidative state as observed by western blot analysis (Fig. 1a). The high reactivity of BAS1 to 10mM H_2_O_2_ compared to the low reactivity of PRXIIE point for the coexistence of two plastids-targeted peroxiredoxin with altered reactivity toward peroxide, possibly providing a mechanism for stepwise sensing of H_2_O_2_ levels, as recently pointed for human peroxiredoxins (Portillo-Ledesma et al., 2018). We also quantified the oxidation response of the catalytic cysteines of the chloroplastic Thioredoxin M2 (TRXM2), Cys110 and Cys113 (ΔOXL=67%, ΔOX_H_ =44%), the catalytic cysteines, Cys^217^ and Cys^220^ involved in the thioredoxin reductase activity of NADPH-dependent thioredoxin reductase C (NTRC, ΔOX_L_=58%), and the catalytic cysteines, Cys^189^ and Cys^192^ of mitochondrial Thioredoxin reductase 1 (NTR1, ΔOX_L_=65%). Reactive Cys were also found in peroxisomal L-ascorbate peroxidase 3 (APX3, Cys^123^, ΔOX_L_=33%), and in GDP-mannose 3,5-epimerase (Cys^223^, ΔOX_L_ =50%, ΔOXH =58%), which is involved in the biosynthesis of L-ascorbate.

**Figure 5.**
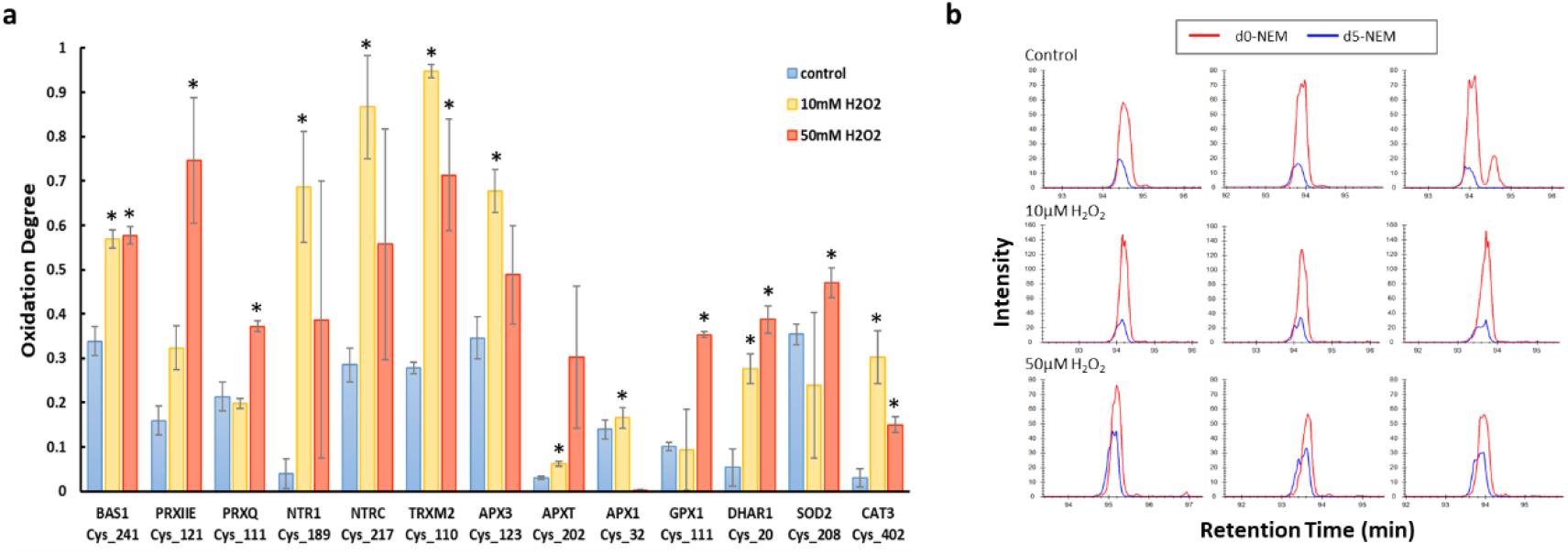
Quantification of cysteine reactivity in redox regulators and antioxidant proteins. **(a)** The oxidation degree of specific Cys residues in several redox regulators and antioxidant proteins. **(b)** Skyline display of the ion chromatograms of the PRXQ peptide (GKPVVLYFYPADETPGCTK), which contains the active site Cys^111^. The ion chromatograms of three independent samples for each treatment are presented in each row, under steady-state and following application of 10mM and 50mM H_2_O_2_.

We also quantified the response of additional antioxidant proteins, such as catalase-3, (CAT3, Cys^402^, ΔOX_L_ =27%, ΔOXH =12%). Cys^145^ and Cys^208^ in copper/zinc superoxide dismutase 1 and 2 (CSD1/SOD1 and CSD2/SOD2), respectively, were also found to be redox-sensitive (ΔOX_H_ =5% and 12%, respectively). These two homologous Cys positions form a regulatory intramolecular disulfide (Cys^59^ with Cys^145^ and Cys^119^ with Cys^208^ in SOD1 and SOD2, respectively). Interestingly, these Cys residues, which were found to be regulated by the thioredoxin and glutaredoxin systems, play a role in the stabilization of the structure of SOD1 in human cells (Cys^57^ with Cys^146^ in hSOD1) and in preventing its aggregation (Álvarez-Zaldiernas et al., 2016). Yet, the role of redox regulation of chloroplast-targeted SOD in plants is yet to be explored.

### Redox sensitivity of proteins involved in chloroplast metabolism

Many of the identified redox-sensitive proteins were assigned to metabolic pathways, highlighting the role of redox post-translational modification in dictating the fluxes through these pathways. Mapping of these redox sensitive proteins to metabolic networks was made using the AraCyc and chloroKB tools (Mueller et al., 2003; Gloaguen et al., 2017), highlighting their participation in many essential pathways such as antioxidant homeostasis, starch metabolism, glycolysis, tricarboxylic acid cycle, the pentose phosphate pathway and photosynthesis (Fig. 6). For example, the extracted ion chromatograms showing the redox sensitivity of Cys^443^ (ΔOX_H_=40%) of chloroplastic glyceraldehyde-3-phosphate dehydrogenase (GAPDH) is presented in Fig. S4. This Cys belongs to the C-terminal extension of the GAPB isoform and is involved in Trx-dependent regulation of GAPDH activity by forming a disulfide bond with Cys^434^, resulting in a kinetically inhibited conformation (Fermani et al., 2007). A similar oxidation response (ΔOX_H_ =34%) was also found for Cys^434^, further demonstrating the sensitivity of these regulatory Cys residues to oxidized conditions (Supp. Table 2). We also observed oxidation of Cys^451^ and Cys^470^ (ΔOX_H_ =19 and 19%, respectively) in the C-terminal extension of RuBisCO activase (RCA), which mediates light regulation of RuBisCO activity (Zhang et al., 2002; Portis Jr et al., 2007). Moreover, we observed oxidation of Cys^449\459^ (ΔOX_L_ =15%) in the large subunit of RubisCO, which are assumed to be involved in redox-dependent inactivation of RubisCO and also contribute to conformational changes that may protect other cysteine residues from further oxidation (Moreno et al., 2008)).

**Figure 6:**
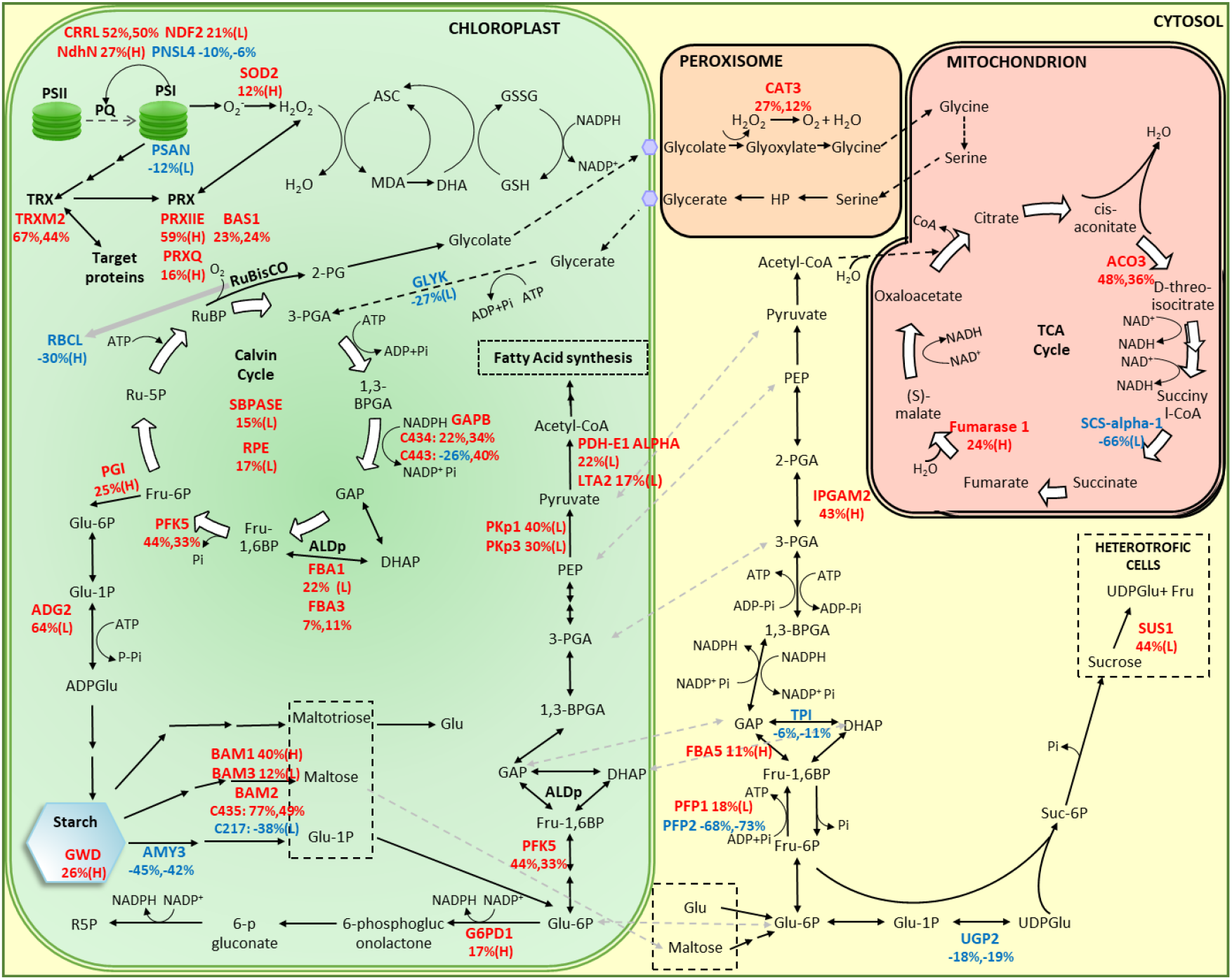
An integrated metabolic map showing redox-sensitive proteins whose oxidative state changed under oxidative stress conditions. Identified redox-sensitive proteins which were oxidized or reduced upon oxidative stress conditions, are shown in red or blue, respectively. Difference in the degree of oxidation (ΔOx) of specific Cys residues in response to 10mM or 50mM H_2_O_2_ as indicated by L (low) and H (high), respectively. A complete list of abbreviations is provided in Table S5. The metabolic map was adopted from Rosa-Téllez et al. (2018), and changes were included to highlight specific metabolic routes.

In addition, we identified several redox-sensitive sites in proteins involved in the cyclic electron flow (CEF). These included Cys^82^ (ΔOX_L_=11%) in PGRL1, which together with PGR5 is involved in antimycin A-sensitive CEF. The identified modified site was found to regulate CEF and NPQ induction by mediating the formation of PGRL1 homodimers through interaction with TRX-m (Hertle et al., 2013; Wolf et al., 2020). Redox regulatory sites were also found in several subunits of NADH dehydrogenase-like (NDH)-complex, including the NDH subunits of subcomplex B2 (Cys^122^, ΔOX_L_=20%) and of subunit U (Cys^106^, ΔOX_L_=51% and ΔOX_H_=50%).

Carbonic anhydrase (CA) catalyzes the reversible interconversion of carbon dioxide (CO_2_) and bicarbonate (HCO_3_ ^−^) and specifically, beta-CAs are highly abundant in plants and diatoms. Reactive Cys residues were identified in Beta carbonic anhydrase 4 (BCA4) (Cys^109^, ΔOX_L_=35%) and Beta carbonic anhydrase 2 (BCA2) (Cys^272^, ΔOX_H_=14%). Redox regulation of CA has been demonstrated in the diatom *Phaeodactylum tricornutum* (Kikutani et al., 2012); however, the role of CA redox regulation in plants was not explored.

We also quantified the oxidation degree of Cys residues that coordinate PSI iron-sulfur clusters such as Cys^573^ in photosystem I P700 chlorophyll a apoprotein A1 (psaA), Cys^559^ in photosystem I P700 chlorophyll a apoprotein A2 (psaB), and Cys^48^ and Cys^54^ in the photosystem I iron-sulfur center (psaC). These Cys undergo only minor oxidation in response to exogenous H_2_O_2_ application. Indeed, O∙^−2^ rather than H_2_O_2_ was shown to inactivate PSI iron-sulfur (FeS) clusters, resulting in PSI photoinhibition under stress conditions (Sonoike et al., 1995; Erling Tjus et al., 1998). As PSI photoinhibition results from photooxidation-induced damage of the iron-sulfur (Fe-S) clusters, the ability to biochemically quantify the redox-state of Cys involved in the coordination of PSI electron carriers, provides a new approach for estimating the PSI state.

We found many redox-sensitive Cys residues involved in starch metabolism (Fig. 6). This included redox regulatory sites in ADP-glucose pyrophosphorylase large subunit 1 (ADG2) and in plastid-localized β-amylases (BAM1-3). For example, we detected significant oxidation of Cys^454^ of BAM1, which was reported to be regulated by TRXs (Valerio et al., 2011). We couldn’t detect the redox state of Cys^82^ of the small subunit of *Arabidopsis* AGPase (APS1), a known redox-regulated site involved in redox activation of starch biosynthesis in the light and in response to sucrose accumulation (Fu et al., 1998). Interestingly, we detected a reduction of Cys^363^ of chloroplastic alpha-amylase 3 (AMY3, ΔOX_L_=−44% and ΔOX_H_=−42%). Notably, a previous investigation of this regulatory site demonstrated that the C363S mutation displayed higher activity, suggesting that reduced Cys at this site is essential for protein function. As reduced AMY3 has been implicated in enzyme activity, our data suggest that starch redox regulation under oxidative stress conditions will induce starch degradation. Since photosynthesis, the process by which plants generate energy-rich organic compounds, may be limited during oxidative stress, inactivation of a starch synthesis enzyme and activation of a starch degradation enzyme can offer alternative energy and carbon sources essential for plant survival.

### Concluding remarks

The SPEAR workflow provides a global system-level quantitative measurement of the thiol-proteome by deferentially labeling reduced and oxidized thiol groups using light and heavy forms of NEM, without enriching labeled peptides. SPEAR enabled simultaneous retrieval of the precise oxidation degree of thousands of Cys sites and protein abundance information from the same MS run. Thus, SPEAR provides a simple, straightforward, and cost-effective approach for studying redox proteomes dynamics. By applying SPEAR to *Arabidopsis* leaves exposed to H_2_O_2_, we offer a comprehensive view of Cys reactivity under oxidative stress conditions. Our dataset contained well-known redox sensors and redox-regulated active sites, which validate the approach, alongside many novel redox-sensitive sites whose roles in plant acclimation to stress conditions remain to be further explored.

## Methods

### Plant growth conditions and treatment

*Arabidopsis thaliana* ecotype Col-0 plants were grown in a greenhouse under a 16/8 h light/dark cycle. Immunoassay, redox imaging and thiol labeling experiments were performed on 3-week-old whole plants, which were incubated in 40ml DW, 10mM H_2_O_2_ or 50mM H_2_O_2_ for 20min at room temperature. Additionally, for redox imaging, plants were incubated with DTT (100mM) and H_2_O_2_ (500mM).

### Immunoassay

Protein extract was prepared in 2ml tubes using a hand homogenizer, in the presence of 1ml cold 10% trichloroacetic acid (TCA) dissolved in water. Proteins were precipitated for 30min on ice, in the dark, and centrifuged at 14,000 rpm, for 20 min, at 4°C. The pellet was then washed 4 times with 100% cold acetone. Residual acetone was removed and the pellet was resuspended in urea buffer comprised of 8M urea, 100mM 4-(2-hydroxyethyl)-1-piperazineethanesulfonic acid (HEPES) (pH 7.2), 1mM EDTA, 2% (w/v) SDS, protease inhibitors cocktail (PI) (Calbiochem) and 100mM N-ethylmaleimide (NEM) (E3876, Sigma) dissolved in EtOH. The samples were incubated for 30min at room temperature and then centrifuged (14,000rpm, 20 min, 4°C) and washed 4 times with 100% cold acetone. The dry pellets were resuspended in urea buffer without NEM and then fractionated on a *4–15*% precast polyacrylamide gel without reducing agent. Sample buffer (x3) was comprised of 150mM Tris-HCl, pH 6.8, 6% (w/v) SDS, 30% glycerol and 0.3% pyronin Y. Fractionated proteins were transferred onto polyvinylidene fluoride membrane (Bio-Rad), using a Trans-Blot Turbo Transfer System (Bio-Rad) with Trans-Blot Turbo Midi Transfer Packs. The membrane was incubated with anti-PRX antibodies (1:1,000) (was kindly provided by Prof. Avichai Danon) or anti-peroxiredoxin-SO_3_ antibody (ab16830) (1:2000), followed by secondary anti-rabbit horseradish peroxidase (1:20,000) (Agrisera), and later developed using standard protocols.

### Redox imaging

Transgenic *Arabidopsis* lines expressing the roGFP probe in specific organelles, including the cytoplasm (Cy-GRX-roGFP2, Marty *et al.* <u>2009</u>), chloroplasts (Chl-roGFP2, with transketolase target peptide (Schwarzlander *et al.* <u>2008</u>), peroxisomes Per-GRX-roGFP2 with peroxisomal targeting signal type 1 SKL amino acid motif, Rosenwasser *et al.* <u>2011</u>) and mitochondria (Mit-roGFP2 with the βATPase signal sequence, Rosenwasser *et al.* <u>2010</u>) were used to map oxidation in response to various redox conditions. Whole-plant roGFP fluorescence images were taken using an in vivo spectral imager (Ami, Spectral Instruments Imaging), following excitation at 465nm (reduction) and 405nm (oxidation) (emission for both: 510nm). Chlorophyll fluorescence was measured following excitation at 405 (emission: 670nm). RoGFP ratiometric images (405/465) were calculated using MATLAB.

### Thiol-labeling assay

Protein extract was prepared in 2ml tubes using a hand homogenizer, in the presence of 1ml cold 20% TCA dissolved in water. Proteins were precipitated for 30min, on ice, in the dark, and centrifuged at 14,000rpm for 20 min, at 4°C. The pellet was washed once with 10% TCA (dissolved in water) and 2 times with 100% cold acetone. The residual acetone was removed by brief evaporation and the precipitate was resuspended (600μl) in urea buffer containing 100mM N-ethylmaleimide (d0-NEM) (E3876, Sigma), dissolved in EtOH, to label the reduced cysteines. Samples were incubated at 50°C for 30min, with 1000rpm shaking. The reaction was terminated and proteins were precipitated by bringing the TCA content to 20% (30min, on ice, in the dark), washed once with 10% TCA and twice with 100% acetone. Samples were then resuspended in urea buffer supplemented with 5.5mM tris 2-carboxyethyl phosphine (TCEP, Sigma) but without NEM, and shaken (1200rpm, 50°C, 30min). Samples were further alkylated by adding 5mM (final concentration) d5-NEM (DLM-6711, CIL) and incubated (50°C, 30min, 1200rpm shaking). TCA (33%) was added, samples were precipitated for 30min, on ice, in the dark and then centrifuged (14,000g, 4°C, 20 min). The pellet was then washed once with 10% TCA and twice with 100% acetone. The acetone was removed by 5min evaporation and samples were stored at −80°C.

An additional set of samples were subjected to thiol labeling as described above, however unlike the former labeling, the reduced cysteines were labeled with d_5_-NEM. Following precipitation with 20% TCA, the samples were resuspended (250μl) in urea buffer containing 100mM d5-NEM (dissolved in EtOH), reduced with 5.5Mm TCEP and further alkylated by adding 5mM (final concentration) d0NEM. The samples were stored at 80°C.

### Sample preparation for LC-MS/MS analysis

All chemicals were from Sigma Aldrich, unless otherwise indicated. Alkylated proteins were suspended in lysis buffer (5% SDS, 50 mM Tris-HCl, pH 7.4). After determination of protein concentration using the BCA assay (Thermo Scientific, USA), 100 μg total protein of each sample was loaded onto S-Trap mini-columns (Protifi, USA), according to the manufacturer’s instructions [1,2]. In brief, after loading, samples were washed with 90:10% methanol/50mM ammonium bicarbonate and then digested (1.5 h, 47 °C) with trypsin (1:50 trypsin/protein). The digested peptides were eluted using 50 mM ammonium bicarbonate and then incubated with trypsin overnight at 37 °C. Two more elution rounds were performed using 0.2% formic acid and 0.2% formic acid in 50% acetonitrile. The three eluted fractions were pooled and vacuum-centrifuged to dryness. Samples were stored at −80 °C until analysis.

### Liquid chromatography

ULC/MS-grade solvents were used for all chromatography steps. Each sample was fractionated offline using high-pH reversed-phase separation, followed by online low-pH reversed-phase separation. Digested protein (100μg) was loaded using high-performance liquid chromatography (Agilent 1260 uHPLC). Mobile phase was: A) 20mM ammonium formate pH 10.0, B) acetonitrile. Peptides were separated on an XBridge C18 column (3×100mm, Waters) using the following gradient: 3% B for 2 min, linear gradient to 40% B over 50min, 5 min to 95% B, maintained at 95% B for 5 min and then back to the initial conditions. Peptides were fractionated into 15 fractions. The following fractions were then pooled: 1 with 8, 2 with 9, 3 with 10, 4 with 11, 5 with 12, 6 with 13 and 7 with 14-15. Each fraction was dried in a speedvac, then reconstituted in 25 μL acetonitrile: water+0.1% formic acid (97:3). Each pooled fraction was then loaded and analyzed using split-less nano-ultra performance liquid chromatography (10 kpsi nanoAcquity; Waters, Milford, MA, USA). The mobile phase was: A) H_2_O + 0.1% formic acid and B) acetonitrile + 0.1% formic acid. Samples were desalted online using a Symmetry C18 reversed-phase trapping column (180μm internal diameter, 20mm length, 5μm particle size; Waters). The peptides were then separated using a T3 HSS nano-column (75μm internal diameter, 250mm length, 1.8μm particle size; Waters) at 0.35 μL/min. Peptides were eluted from the column into the mass spectrometer, using the following gradient: 4% to 27% B over 105 min, 27% to 90%B over 5 min, maintenance at 90% for 5 min and then back to the initial conditions.

### Mass spectrometry

The nanoUPLC was coupled online through a nanoESI emitter (10 μm tip; New Objective; Woburn, MA, USA) to a quadrupole orbitrap mass spectrometer (Orbitrap Fusion Lumos, Thermo Scientific) using a FlexIon nanospray apparatus (Proxeon). Data were acquired in DDA mode, using a 3-second cycle method. MS1 was performed in the Orbitrap with resolution set to 120,000 (at 200m/z) and maximum injection time set to 50msec. MS2 was performed in the Orbitrap after HCD fragmentation, with resolution set to 15,000 and maximum injection time set to 100msec.

### Data analysis - protein expression

Raw data were analyzed using MaxQuant V1.6.0.16.[3]. MS/MS spectra were searched against the *Arabidopsis thaliana* protein database downloaded from UniProt (www.Uniprot.org). Enzyme specificity was set to trypsin and up to two missed cleavages were allowed. Variable modifications were set to oxidation of methionines, and light (d0) or heavy (d5) NEM labeling of cysteines (+125.0477 or +130.0791 Da, respectively). Peptide precursor ions with a maximum mass deviation of 4.5 ppm and fragment ions with a maximum mass deviation of 20 ppm were searched. Peptide, protein and site identifications were filtered at an FDR of 1%, using the decoy database strategy. The minimal peptide length was 7 amino acids and the minimum Andromeda score for modified peptides was 40. Peptide identifications were propagated across samples using the match-between-runs option. Searches were performed with the label-free quantification option selected. Fold changes were calculated based on the ratio of geometric means of the case versus control samples. A Student’s t-test, after logarithmic transformation, was used to identify significant differences across the biological replica.

### Data analysis – Cys oxidation

Peptide identification and quantification were conducted using the MetaMorpheus software (v0.0.297 https://smith-chem-wisc.github.io/MetaMorpheus/ or https://github.com/smith-chem-wisc/MetaMorpheus). RAW MS files were searched against the *Arabidopsis thaliana* (Mouse-ear cress) proteome downloaded from UniProt (UniProtKb Proteome ID UP000006548, as of December 14, 2000, in XML format). All MS files, including the sample in which reduced thiols were labeled with d0-NEM and those labeled with d5-NEM, were jointly processed. In short, the MetaMorpheus calibration task and the G-PTM-D task were performed using the default parameters, with the following modifications: nethylmaleimide on C and Nem:2H(5) on C were set as G-PTM-D modifications. The following options were checked in the search settings: apply protein parsimony and construct protein groups, treat modified peptides as different peptides, match between runs, normalize quantification results. An additional MetaMorpheus run was conducted without the G-PTM-D algorithm, using Nethylmaleimide on C and Nem:2H(5) on C as variable modifications in the search task. Detailed MetaMorpheus parameters used for both runs are provided in Table S4.

The generated output files AllQuantifiedPeptides.tsv were used for further analysis. First, files were trimmed, and only cysteine-containing peptides labeled with d0 or d5-NEM (belongs to the core set in which reduced Cys was labeled with d0-NEM) were selected. Next, a list with peptides bearing both labels was extracted (pairs). A CysID was then assigned to each cysteine-containing peptide; the ID included the UniProt identifier name of the protein and the NEM-labeled cysteine residue position in the protein sequence. Next, the intensity of each peptide throughout the six fractions was summed. If a specific Cys was represented by more than one peptide, the overall intensity of peptides with identical CysIDs was examined and the peptide with the highest intensity was selected for further analysis. The oxidation degree (OxD) of a Cys was calculated using the following formula: 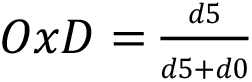, where d5 represents the intensity of the d5-NEM labeled peptide and d0 the intensity of the d0-NEM labeled peptide. Only peptides identified in at least two out of three biological replicates were included. Delta oxidation (ΔOx) was calculated by subtracting the OxD of H_2_O_2_-treated plants from the OxD of control plants. The p-value for each peptide was determined using student’s t-test.

The OxD for peptides with 2 Cys was calculated twice using the formulas: 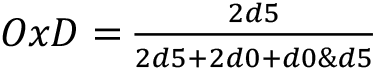 or 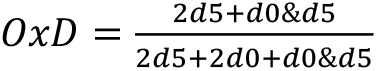, where 2d5 represents the intensity of the peptide with 2 Cys that both Cys were d5-NEM labeled, 2d0 the intensity of the peptide with 2 Cys that both Cys were d0-NEM labeled, and d0&d5 the intensity of the peptide with 2 Cys that one Cys was d0-NEM labeled and the other Cys was d5-NEM labeled. The two corresponding ΔOx values were calculated and the p-value for each peptide was determined using student’s t-test, accordingly.

Only peptides which passed the threshold of ΔOx > |±10%| and α<0.05 (at least in one of the applied H_2_O_2_ concentrations) were considered “redox-sensitive peptides”. Some of the “redox-sensitive peptides” were imported into Skyline (Pino et al., 2020; an open-source program, can be downloaded at http://proteome.gs.washington.edu/software/skyline) for visualization of the d0- and d5-NEM peak pairs.

### Functional annotation analysis

The DAVID (Database for Annotation, Visualization and Integrated Discovery) Bioinformatics Resources tool v6.7was used for mapping protein into Gene Ontology terms (GO) and KEGG (Kyoto Encyclopedia of Genes and Genomes) pathways and for enrichment analysis. Further mapping into metabolic pathways was carried out using the *Arabidopsis* chloroplast knowledge base (cloroKB, Gloaguen et al., 2017), the AraCyc database from the Plant Metabolic Network (PMN, Mueller et al., 2003), and manual annotation. Subcellular protein localization was predicted using the SUBA tool (the central resource for *Arabidopsis* protein subcellular location data). Organelle-specific enrichment for the proteins whose redox state had changed was calculated based on the cumulative distribution function (CDF) of the hypergeometric distribution (https://systems.crump.ucla.edu/hypergeometric/). ΔOx of Cys belonging to various subcellular compartments was presented using the PlotsOfData tool (https://huygens.science.uva.nl/PlotsOfData/). The Probability logo generator for biological sequence motifs (Plogo, https://plogo.uconn.edu/) was used to estimate the probability of the occurrence of specific amino acid sequence motifs around the NEM labeled Cys. The functional annotation regarding the active and binding sites, metal binding and disulfide bonds, for all the quantified cysteines was retrieved from the UniProt Knowledgebase (https://www.uniprot.org/uploadlists/). The enrichment probability for the "redox-sensitive cysteines" compared to all the quantified cysteines was calculated based on the cumulative distribution function (CDF) of the hypergeometric distribution (https://systems.crump.ucla.edu/hypergeometric/).

## Author contributions

S.D., N.L., A.S., Y.L., and S.R. conceived and designed the research. N.L. and A.S., performed the research. S.D., A.S., Y.L and S.R. analyzed the data. S.D. and S.R. wrote the manuscript with inputs from N.L., A.S., and Y.L.

## Acknowledgments

We thank Dana Reichmann for her critical comments on the manuscript.

## Supplemental data

The following is the Supplementary data to this article:

**Figure S1:** RT difference between peptides labeled with d0-NEM and d5-NEM.

**Figure S2:** Consensus motif of identified Cys containing peptides not labeled with NEM.

**Figure S3:** Skyline display of the ion chromatograms of the BAS1 peptide (TLQALQYIQENPDEVCPAGWKPGEK).

**Figure S4:** Skyline display of the ion chromatograms of the GAPB peptide (TNPADEECKVYD).

**Figure S5:** Violin plot visualization of the difference in the thiol-oxidized state between steady state and exposure to H_2_O_2_ treatment in different subcellular compartments.

**Table S1:** A list of 17 significantly redox-sensitive peptides derived from using the regular d0-NEM and d5-NEM labeling of reduced and oxidized Cys thiols, respectively.

**Table S2:** A list of 577 significantly redox-sensitive peptides derived from the combined approach including the regular d0-NEM and d5-NEM labeling and oppositely labeled “marker” samples.

**Table S3:** Changes in proteins expression levels under oxidative stress conditions compared to control.

**Table S4:** MetaMorpheus setting parameters used for the two runs, with and without the G-PTM-D task, referred to as “method 1” and “method 2”, respectively.

**Table S5:** The pathways abbreviations for the integrated metabolic map presented in Fig. 6.

