## Supplemental Figures for "SPEAR: a proteomics approach for simultaneous protein expression and redox analysis"

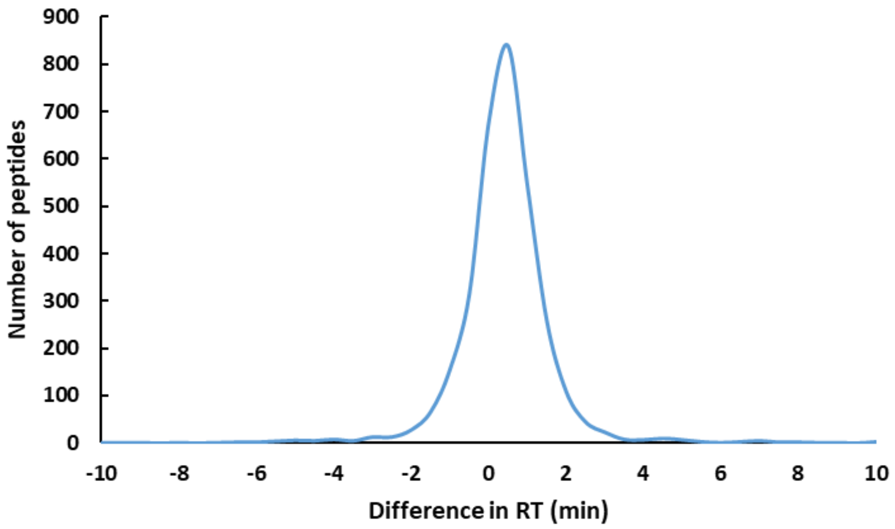


**Figure S1: RT difference between peptides labeled with d0-NEM and d5-NEM.** The differences in RT between peptides with the same sequence labeled with d0-NEM or d5-NEM were calculated for 3173 peptide pairs and presented as distribution.

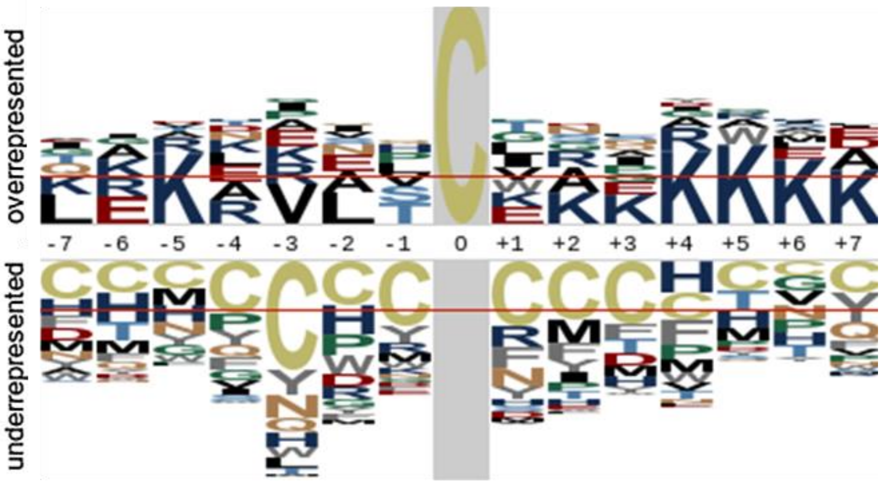


**Figure S2: Consensus motif of identified Cys containing peptides not labeled with NEM.** *Arabidopsis thaliana* proteome was used as the proteomic background population for the calculation of the probabilities.

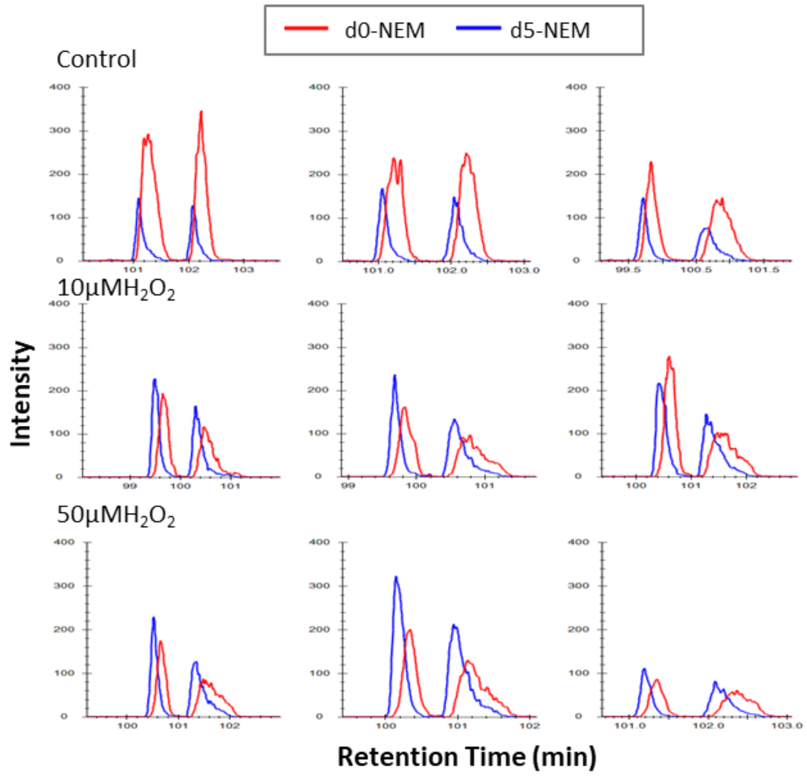


**Figure S3: Skyline display of the ion chromatograms of the BAS1 peptide (TLQALQYIQENPDEVCPAGWKPGEK).** This peptide contains $Cys^{241}$, which forms a disulfide bond with $Cys^{119}$. The ion chromatograms of three independent samples for each treatment are presented in each row, under steady-state and following application of 10mM and 50mM $H_2O_2$.

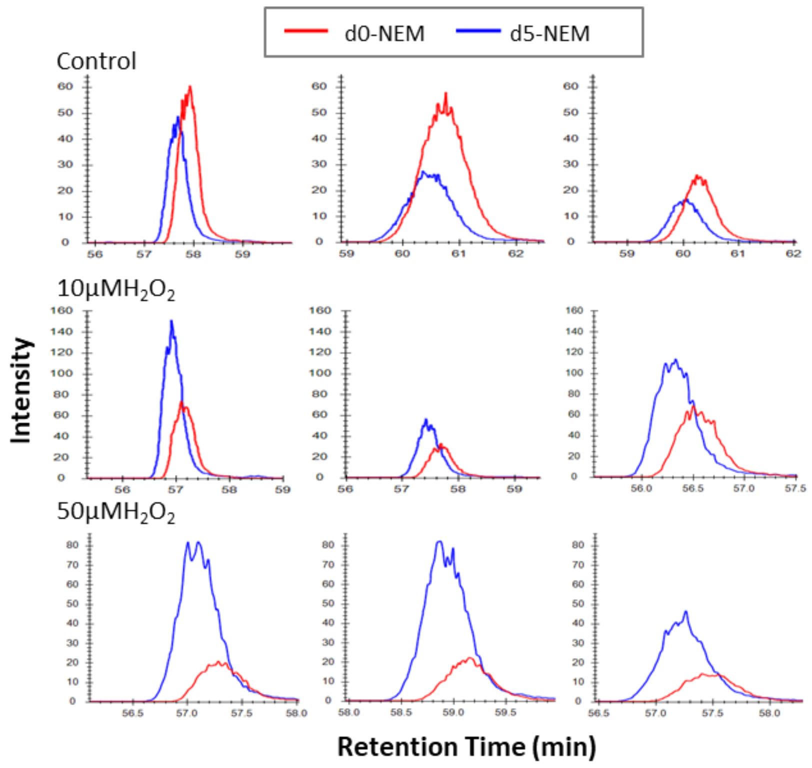


**Figure S4: Skyline display of the ion chromatograms of the GAPB peptide (TNPADEECKVYD).** This peptide contains $Cys^{443}$, which forms a disulfide bond with $Cys^{434}$ (also identified in our data). The ion chromatograms of three independent samples for each treatment are presented in each row, under steady-state and following application of 10mM and 50mM $H_2O_2$.

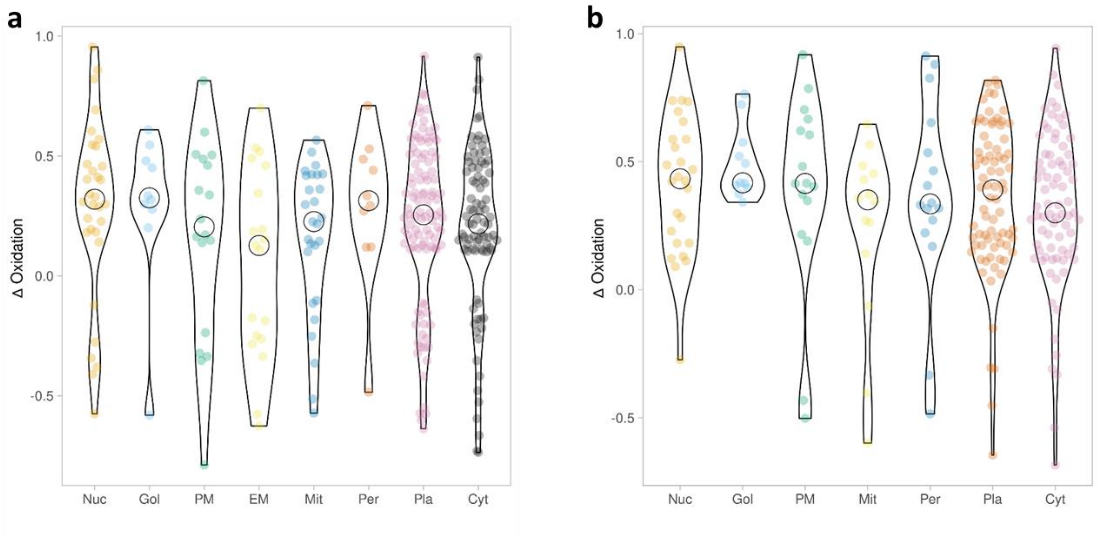


**Figure S5: Violin plot visualization of the difference in the thiol-oxidized state between steady state and exposure to $H_2O_2$ treatment in different subcellular compartments.** Violin plot visualization of the difference in the thiol-oxidized state between steady state and exposure to 50mM **(a)** and 10mM **(b)** $H_2O_2$ treatment. Nuc, nucleus; Gol, golgi; PM, plasma membrane; EM, extracellular matrix; Mit, mitochondrion; Per, peroxisome; Pla, plastid; Cyt, cytosol.
